# Model-based analysis of the circadian rhythm generation of bioluminescence reporter activity in duckweed

**DOI:** 10.1101/2024.05.26.595939

**Authors:** Yu Horikawa, Emiri Watanabe, Shogo Ito, Tokitaka Oyama

## Abstract

Bioluminescence monitoring techniques are widely used to study the gene expression dynamics in living plants. Monitoring the bioluminescence from a luciferase gene under the control of a circadian promoter is indispensable for examining plant circadian systems. The bioluminescence monitoring technique was successfully applied to physiological studies of circadian rhythms in duckweed plants. It has been reported that a luciferase gene under a constitutive promoter also exhibits a bioluminescent circadian rhythm in duckweed. However, the mechanisms underlying rhythm generation remain unknown. In this study, we performed a model-based analysis to evaluate the machinery that generates the bioluminescence rhythm. We hypothesized the rhythmic factor of three aspects regarding the bioluminescence intensities of luciferase in cells: luminescence efficiency, production rate, and degradation rate. Theoretically, if the latter two are involved in rhythm generation, the difference in luciferase stability affects the amplitude and phase relations of the bioluminescence rhythm. Luciferase stability is irrelevant to these rhythm properties if only the luminescence efficiency is involved. First, we simulated the bioluminescence rhythms of two luciferases with different stabilities associated with each of three rhythmic factors. Luciferase stability was set based on the reported values for Emerald-luciferase and Emerald-luciferase-PEST. We then experimentally examined the bioluminescence rhythms of reporters of these luciferases driven by the *CAULIFLOWER MOSAIC VIRUS 35S* promoter in the duckweed *Lemna japonica*. Their circadian properties matched those obtained from the simulation of the luminescence efficiency. This supports the view that cells in duckweed show circadian changes in physiological conditions associated with the luciferase enzyme reaction.

---

Plant physiology is regulated by day/night environmental cycles. With its self-sustain clock, the circadian system is critically involved in diel regulation (Greenham and McClung. 2015). The circadian regulation of gene expression is based on the circadian system. Monitoring gene expression in living organisms is essential for studying circadian systems (Millar et al. 1992). A luciferase reporter with a circadian promoter is widely used to analyze gene expression dynamics due to its non-invasive nature and quantitative performance (Muranaka et al. 2013). Spatiotemporal analysis of the plant circadian system at the single-cell level was performed using a bioluminescence monitoring system in duckweed transfected with a luciferase reporter (Muranaka et al. 2016). It has been reported that the bioluminescence of luciferase driven by a constitutive promoter, such as the *CAULIFLOWER MOSAIC VIRUS 35S* promoter (*CaMV35S*), showed a circadian rhythm in various duckweeds (Lemnaceae) (Muranaka et al. 2015). The mechanism of rhythm generation is unknown; it has been suggested that physiological conditions associated with the luciferase reaction in cells mediate rhythms post-transcriptionally (Watanabe et al. 2021, Watanabe et al. 2023). In this scenario, a bioluminescence rhythm appeared even when the amount of luciferase in the cells was constant. We performed a model-based analysis of the circadian rhythms of the bioluminescence of two luciferases with different stabilities to address how the bioluminescence rhythm is generated. Theoretically, stability is associated with circadian properties, such as amplitude and phase relations, when the production and/or degradation rates of luciferase are rhythmic (Lück et al., 2014). We experimentally evaluated the rhythm generation scenarios by analyzing the bioluminescence rhythms of stable and unstable luciferase.

We used *Lemna japonica* 5512 [*Lemna minor* 5512 in previous studies: the Rutgers Duckweed Stock Cooperative (http://www.ruduckweed.org/)] for the experiments. Duckweed plants were maintained in NF medium containing 1% sucrose under constant light conditions in a temperature-controlled room (25°C), as previously described (Muranaka et al. 2015). We used *pUC18-CaMV35S::ELUC* (*CaMV35S::ELUC*) and *pUC18-CaMV35S::ELUC-PEST* (*CaMV35S::ELUC-PEST*) as circadian bioluminescence reporters. *ELUC* encodes emerald luciferase (ELuc) derived from the Brazilian click beetle (*Pyrearinus termitilluminans*) (TOYOBO; Nakajima et al. 2010). These reporters were modified versions of *pUC18-CaMV35S::LUC+*, as described previously (Muranaka et al. 2013). Transfection of a luciferase reporter using particle bombardment and bioluminescence monitoring of the transfected duckweed plants were performed as described previously (Muranaka et al. 2015). The luminescence of the sample dish was measured every 20 min using a photomultiplier tube. Single-cell bioluminescence imaging was performed using an EM-CCD camera (ImagEM C9100-13; Hamamatsu Photonics) with a camera lens (XENON 0.95/25MM C-mount; Schneider Optics) as previously described (Muranaka and Oyama 2016). Bioluminescence images (16-bit TIFF format) of plants 3 days after gene transfection were captured with the ImagEM camera (cooled at −80ºC) at an EM gain of 1,200 with 3, 5, 15, 30, and 60-s exposures after at least 4 min dark treatment for autofluorescence decay. Time series analysis was implemented using Python 3.8.8 (NumPy 1.20.1, SciPy 1.6.2). The data analysis codes are available at https://github.com/ukikusa/circadian_analysis v0.9. Further code and data are available upon request from the corresponding authors.

We first performed a model-based analysis to identify the rhythmic factor(s) that generate the bioluminescence rhythm of a luciferase driven by a constitutive promoter. We focused on the following three aspects regarding the bioluminescence intensities of cells expressing the luciferase enzyme: luminescence efficiency, production rate, and degradation rate. The structure of the bioluminescence system is shown in Figure 1A, luminescence emission (*L*(*t*)) is the output, and the luciferase amount (*x*(*t*)) is the state variable. A circadian rhythm in luminescence efficiency can confer rhythmicity to bioluminescence even when the luciferase amount is constant. A circadian rhythm in the production or degradation rate results in a circadian rhythm in luciferase amounts, which consequently generates a bioluminescence rhythm. The stability of the luciferase enzyme can affect the phase and amplitude of the bioluminescence rhythm when the amount of luciferase is associated with the rhythm (Lück et al., 2014). We analyzed three simple cases in a simulation model for the bioluminescence system to compare the rhythmic outputs of stable luciferase (StLUC) and unstable luciferase (UnstLUC). We set two luciferases with degradation rates *γ*_StLUC_ = 0.07 h^−1^ and *γ*_UnstLUC_ = 0.23 h^−1^. These values for *γ*_StLUC_ and *γ*_UnstLUC_ correspond to those of ELUC (half-life: 10 h) and ELUC-PEST (half-life: 3 h), respectively (Yasunaga et al. 2015). ELUC-PEST is a short-life-type ELUC containing a 40-aa PEST sequence at the C-terminus. The protein levels of these two luciferases differed 3.3-fold in their steady states (Figure 1B; SIM B in Table 1). If the rhythmic factor was simply set to the luminescence efficiency, the amplitudes of the bioluminescence rhythms were different, but the relative values of StLUC and UnstLUC were the same (Figure 1C; SIM C in Table 1). In this case, the phases are the same. If the rhythmic factor is set to the production rate, the amplitudes of the bioluminescence rhythms are similar between the two luciferases (Figure 1D, left; SIM D in Table 1). Regarding the amplitudes of the relative values, UnstLUC showed a larger magnitude than StLUC (Figure 1D, right; SIM D in Table 1). A larger phase delay occurred in the StLUC rhythm (SIM D in Table 1). As shown in Figure 1E and Table 1 (SIM E), the rhythmic factor of the degradation rate resulted in circadian rhythms for both luciferases similar to those generated by the rhythmic production rate.

**Table 1.** Properties of circadian rhythms in each simulation and the experimental results.

| Analysis | Luminescence intensities<sup>a</sup> |  |  | Relative amplitude |  |  | Phase property [h] |  |  |
| --- | --- | --- | --- | --- | --- | --- | --- | --- | --- |
|  | StLUC | UnstLUC | St / Unst | StLUC | UnstLUC | St / Unst | StLUC | UnstLUC | St – Unst |
| SIM B | 14.28 | 4.35 | 3.28 | NA | NA | NA | NA | NA | NA |
| SIM C | 14.28 | 4.35 | 3.28 | 0.22 | 0.22 | 1.00 | 0 | 0 | 0 |
| SIM D | 14.28 | 4.35 | 3.28 | 0.06 | 0.15 | 0.41 | –5.24 | –3.27 | –1.97 |
| SIM E | 14.31 | 4.41 | 3.24 | 0.06 | 0.15 | 0.41 | 7.77 | 9.54 | –1.77 |
| EXP<sup>b</sup> | 84.60 | 18.24 | 4.64 | 0.18 ± 0.03 | 0.22 ± 0.04 | 0.83 | 0<sup>c</sup> | 0.78<sup>c</sup> | –0.78 |

<sup>a</sup>For simulation data, arithmetic means of luminescence intensities in the period between 93 h and 119 h (Figure 1). For experimental data, geometric means of cellular bioluminescence intensities (photons min<sup>–1</sup>) at a time point (Figure 2A). <sup>b</sup>Experimental data shown in Figure 2A and 2C are summarized for StLUC (ELUC) and UnstLUC (ELUC-PEST). <sup>c</sup>The mean time at the third trough for StLUC is set to 0 h.

**Figure 1.**
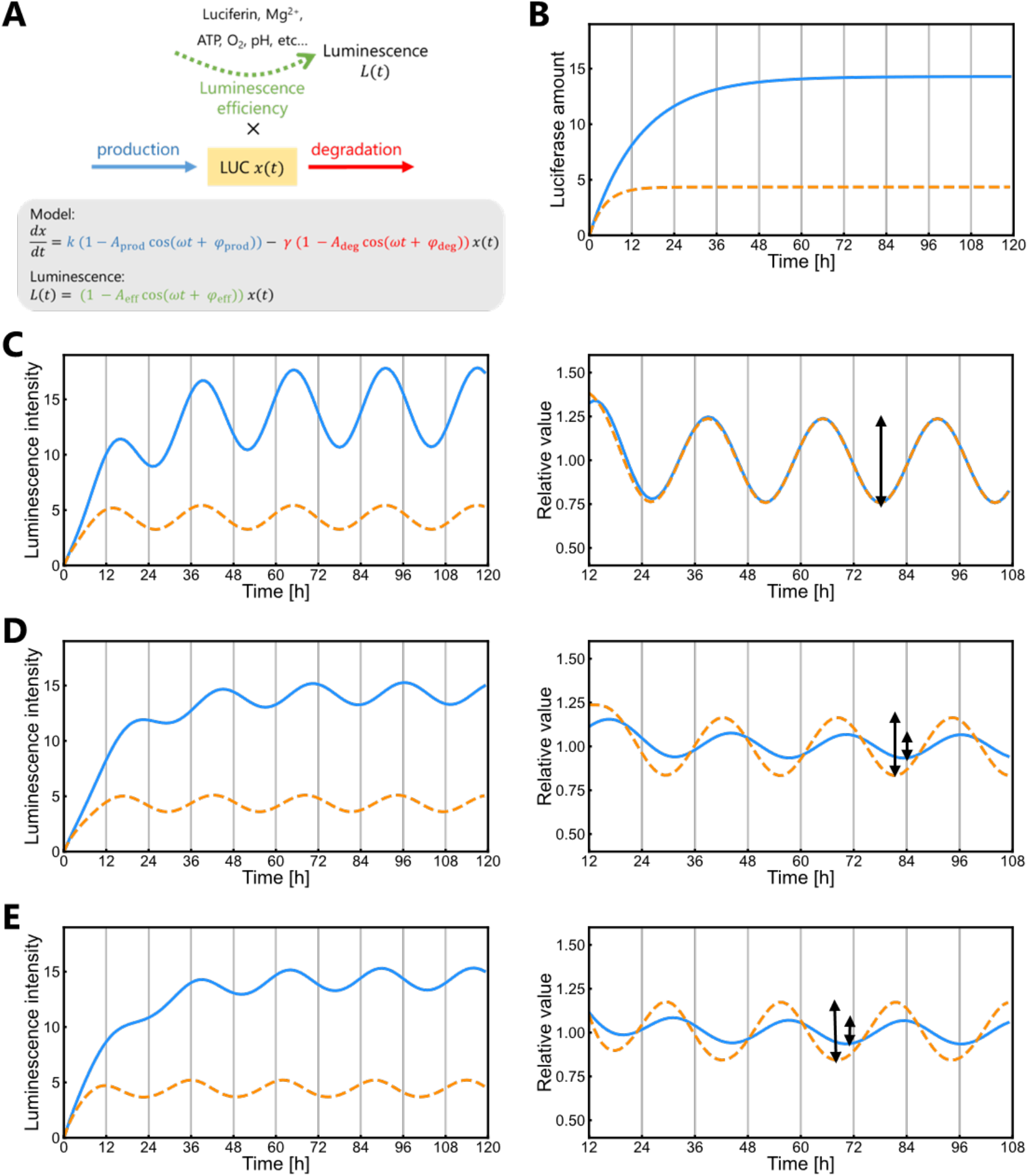
Simulation of bioluminescence behavior in the three conditions for rhythmic factor (A) Model. The amount of LUC (*x*(*t*)) is the state variable and luminescence intensity (*L*(*t*)) is the output. The production rate coefficient, the degradation rate coefficient, and the coefficient of luminescence efficiency are marked in blue, red, and green, respectively. *k*and *γ* are the mean rate coefficients. Time modulation of *x*(*t*) and *L*(*t*) is described by cosine-shaped functions with relative amplitudes *A*_prod_, *A*_deg_,*andA*_eff_ (values between 0 and 1). (B-E) Simulation of the model. Comon parameter values: *ω =* 2π/26, *φ*_prod_ *= φ*_eff_ *= φ*_eff_ *=* 0, *k=* 1 h^−1^, *γ*_UnstLUC_ *=* 0.07 h^−1^. and *γ*_UnstLUC_ *=* 0.23 h^−1^. A solid blue line and a dashed orange line indicate StLUC and UnstLUC, respectively. Simulation of the luciferase amount without the rhythm in the production rate and degradation rate (SIM B; *A*_prod_ *= A*_deg_ *= A*_eff_ *=* 0) is shown in (B). Simulation of luminescence intensities and relative values with a circadian rhythm in the luminescence efficiency (SIM C; *A*_eff_ *=* 0.25, *A*_prod_ *= A*_deg_ *=* 0) (C), the production rate (SIM D; *A*_prod_ *=* 0.25, *A*_deg_ *= A*_eff_ *=* 0) (D), and the degradation rate (SIM E; *A*_deg_ *=* 0.25, *A*_prod_ *= A*_eff_ *=* 0) (E). The relative values are normalized by the 26-h moving average at each time point. The two-headed arrow for a rhythm represents twice the amplitude and the point of the third trough.

Next, we compared the bioluminescence rhythms of *CaMV35S::ELUC* (*ELUC*) and *CaMV35S::ELUC-PEST* (*ELUC-PEST*) in cells. The bioluminescence intensities of individual cells were measured using single-cell bioluminescence imaging (Figure 2A). The intensity values of both the luciferase reporters showed a log-normal distribution. The geometric mean for *ELUC* was 4.6-fold as large as that for *ELUC-PEST*. This difference was comparable to the simulation results (Table 1). Bioluminescence for both luciferases was monitored under constant light (Figure 2B). The bioluminescence intensities of both reporters exhibited circadian rhythms. While the bioluminescence intensities of *ELUC-PEST* gradually decreased, the circadian rhythms of relative values matched those of *ELUC* (Figure 2B and 2C; EXP in Table 1). A phase difference was observed between the two reporters in the third trough (Figure 2C; EXP in Table 1). However, the first trough time was almost the same for *ELUC* and *ELUC-PEST*, suggesting that these two luciferase reporters showed a similar phase relation (Figure 2C). In summary, matching of the bioluminescence rhythms of relative values with a similar phase was a characteristic of the SIM C results (Figure C; Table 1). Thus, we concluded that the circadian rhythm of luciferase expressed under the control of the *CaMV35S* promoter was generated at the step of luminescence efficiency for the luciferase enzyme reaction. This conclusion does not exclude the possibility that a circadian rhythm in the production rate or degradation rate may result in a bioluminescence rhythm. However, its magnitude should be much smaller than that of the luminescence efficiency.

**Figure 2.**
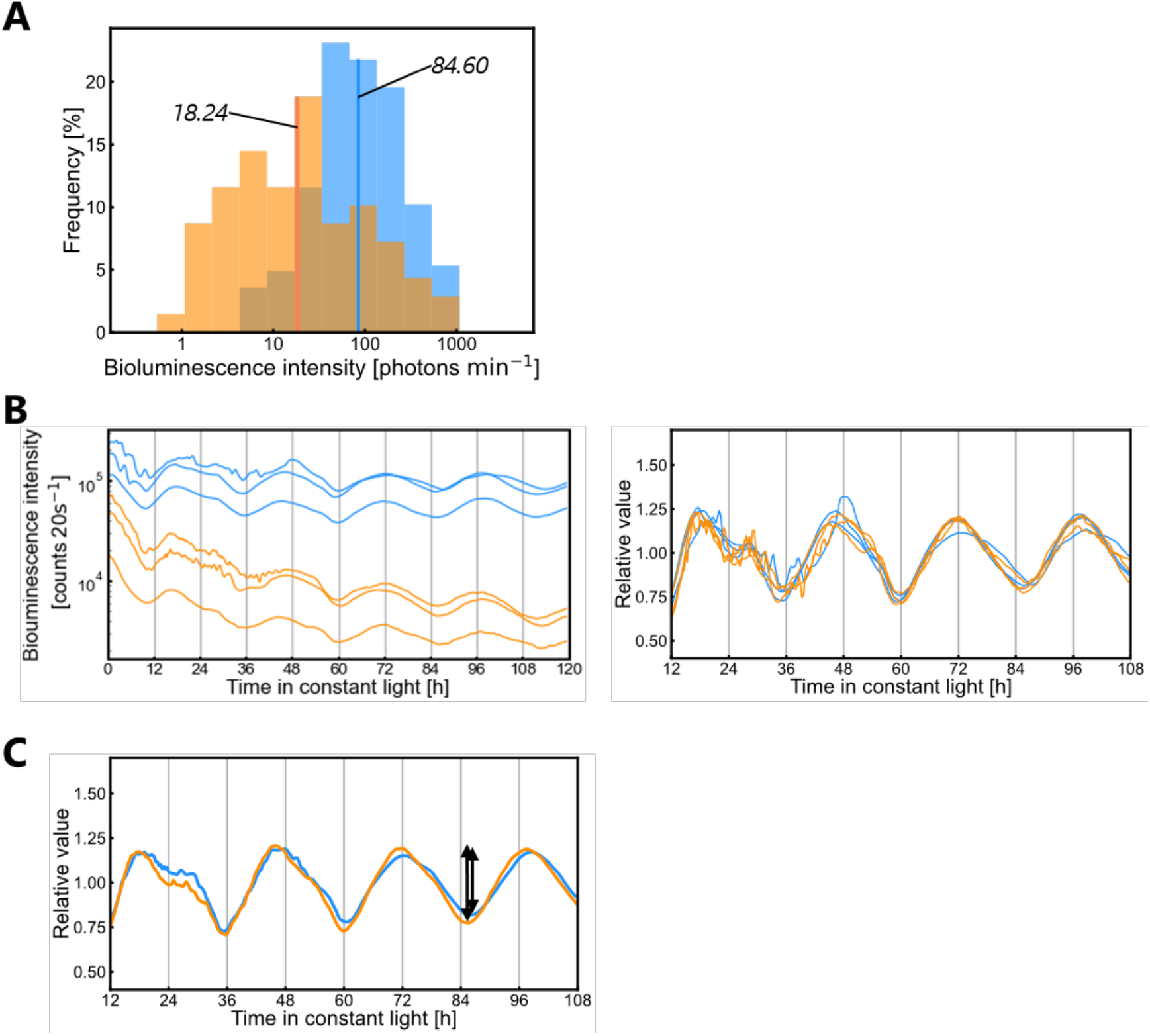
Bioluminescence of *CaMV35S::ELUC* and *CaMV35S::ELUC-PEST* (A) Frequency distributions for cellular bioluminescence intensities of *ELUC* (blue) and *ELUC-PEST* (orange). Geometric means are indicated in the graph. Both histograms show a log-normal distribution. (Kolmogorov–Smirnov test, *P* > 0.05; *n* = 225 (*ELUC*), *n* = 69 (*ELUC-PEST*)). Note that the x-axis is on a logarithmic scale. (B) Circadian rhythms of bioluminescence intensities (left) and relative values (right) for *ELUC* (blue lines) and *ELUC-PEST* (orange lines) under constant light conditions. Relative values are the normalized values by the 26-h moving average at each time point. Plants subjected to gene transfection were entrained to two 12-h dark/12-h light cycles and then released into constant light conditions for bioluminescence monitoring using a photomultiplier tube. Time-series data of three replicates in an experiment among three independent experiments are shown. (C) Means of relative values of all nine samples in the three experiments are shown. A two-headed arrow for a rhythm represents twice the amplitude and the point of the third trough.

The mechanism for generating the circadian rhythm at the luminescence efficiency step is unknown. However, it is clear that the physiological conditions in the cytoplasm associated with the luciferase reaction in duckweed show a circadian rhythm. The bioluminescence of any luciferase reporter is affected by circadian changes in the luminescence efficiency. Various luciferase reporters driven by a circadian promoter at a specific phase have been used to study circadian rhythms in duckweed (Serikawa et al. 2008, Isoda et al. 2022, Watanabe et al. 2023). The bioluminescence rhythms of these reporters include at least two circadian components: the production rate and luminescence efficiency. Therefore, it is necessary to remove the circadian component of luminescence efficiency to analyze the circadian behavior of promoter activity.

## Acknowledgements

This work was supported in part by the Japan Society for the Promotion of Science KAKENHI [grant numbers JP21J23250 (E.W.), JP20K06342 (S.I.), 24H02121 (S.I.), 17KT0022 (T.O.), JP19H03245 (T.O.)], the Japan Science and Technology Agency (JST), JST ALCA (JPMJAL1108, T.O.), and JST SATREPS (JPMJSA2004, S.I., and T.O.). We thank S. Honda and R. Ueno for their support with single-cell bioluminescence imaging. We also thank Dr. Masaaki Morikawa for providing us with *L. japonica* 5512.

